# Relationship between simultaneously recorded spiking activity and fluorescence signal in GCaMP6 transgenic mice

**DOI:** 10.1101/788802

**Authors:** Lawrence Huang, Ulf Knoblich, Peter Ledochowitsch, Jérôme Lecoq, R. Clay Reid, Saskia E. J. de Vries, Michael A. Buice, Gabe J. Murphy, Jack Waters, Christof Koch, Hongkui Zeng, Lu Li

## Abstract

Two-photon calcium imaging is often used with genetically encoded calcium indicators (GECIs) to investigate neural dynamics, but the relationship between fluorescence and action potentials (spikes) remains unclear. Pioneering work linked electrophysiology and calcium imaging *in vivo* with viral GECI expression, albeit in a small number of cells. Here we characterized the spike-fluorescence transfer function *in vivo* of 91 layer 2/3 pyramidal neurons in primary visual cortex in four transgenic mouse lines expressing GCaMP6s or GCaMP6f. We found that GCaMP6s cells have spike-triggered fluorescence responses of larger amplitude, lower variability and greater single-spike detectability than GCaMP6f. Mean single-spike detection rates at high spatiotemporal resolution measured in our data was >70% for GCaMP6s and ~40-50% for GCaMP6f (at 5% false positive rate). These rates are estimated to decrease to 25-35% for GCaMP6f under generally used population imaging conditions. Our ground-truth dataset thus supports more refined inference of neuronal activity from calcium imaging.

## Introduction

Genetically encoded calcium indicators (GECIs) are widely used with two-photon (2-p) laser scanning microscopy to report neuronal activity within local populations *in vivo* (Luo et al., 2018). This optical approach is minimally invasive and enables simultaneous measurement of activity from hundreds or even thousands of neurons at single-cell resolution, over multiple sessions. Using a contemporary GECI such as GCaMP6s, fluorescence changes associated with isolated single spikes (action potentials) *in vivo* can be detected when imaged at sufficiently high spatiotemporal resolution (Chen et al., 2013) (http://dx.doi.org/10.6080/K02R3PMN). However, despite recent advances in imaging approaches and GECI development, calcium imaging remains an indirect measure of a neuron’s spiking activity. Inferring the underlying spike train or firing rate from calcium imaging remains challenging (Theis et al., 2016; Berens et al., 2018), because the spike to calcium-dependent fluorescence transfer function may be different for each neuron due to a variety of intrinsic and extrinsic factors. Thus, investigating the direct relationship between spiking activity and calcium-based fluorescence signal in a large number of neurons by simultaneous fluorescent imaging and electrical recording *in vivo* can facilitate better understanding of neuronal activities through calcium imaging studies.

Compared to viral expression, transgenic mouse lines offer convenience (e.g. bypassing virus injection and associated procedures) and achieve more uniform GECI expression in genetically defined neuronal populations (Madisen et al., 2015; Daigle et al., 2018). We might expect the spike to calcium-dependent fluorescence transfer function to be relatively uniform across neurons and animals. Using our intersectional transgenic mouse lines that enable Cre recombinase-dependent expression of GCaMP6s or GCaMP6f, we simultaneously characterized the spiking activity and fluorescence of individual GECI-expressing pyramidal neurons in layer (L) 2/3 of mouse primary visual cortex (V1). Consisting of 91 neurons from 4 mainstream transgenic lines, this ground truth dataset provides quantitative insight into the relationship between *in vivo* spiking activity and observed fluorescence signals, and will aid the interpretation of existing and future calcium imaging datasets.

## Results

### Dataset overview

To characterize the single-cell transfer function between observed fluorescence signals and underlying spikes *in vivo*, we performed simultaneous calcium imaging and cell-attached recordings in V1 L2/3 excitatory pyramidal neurons in anesthetized mice (**Figure 1A**, see Methods). The mice were from 4 transgenic lines, including 2 GCaMP6s-expressing lines, *Emx1* -IRES-Cre;*Camk2a*-tTA;Ai94 (referred to as Emx1-s in this study) and *Camk2a-tTA;tetO-* GCaMP6s (tetO-s), and 2 GCaMP6f-expressing lines, *Emx1* -IRES-Cre;*Camk2a*-tTA;Ai93 (Emx1-f) and *Cux2*-CreERT2;*Camk2a*-tTA;Ai93 (Cux2-f) (**Table 1**). A total of 237 neurons, all with fluorescence excluded from the nucleus, were randomly selected to record and image for spontaneous activity or visually-evoked responses. To directly compare our results to virally-expressed GCaMP6f and GCaMP6s (Chen et al., 2013) (http://dx.doi.org/10.6080/K02R3PMN), calcium imaging was performed at high optical zoom focused on individual cells, with a field of view (FOV) of ~20 x 20 μm and scanning rate at ~158 frames per second (fps). We also patched and imaged a subset of these neurons at a lower zoom factor, *i.e.* one at which the responses of many neurons can be characterized in parallel (FOV of ~400 x 400 μm at ~30 fps), allowing direct comparison of calcium fluorescence responses acquired at higher spatiotemporal resolution with that more commonly employed in typical calcium imaging studies including our Brain Observatory dataset (de Vries et al., 2019).

**Figure 1.**
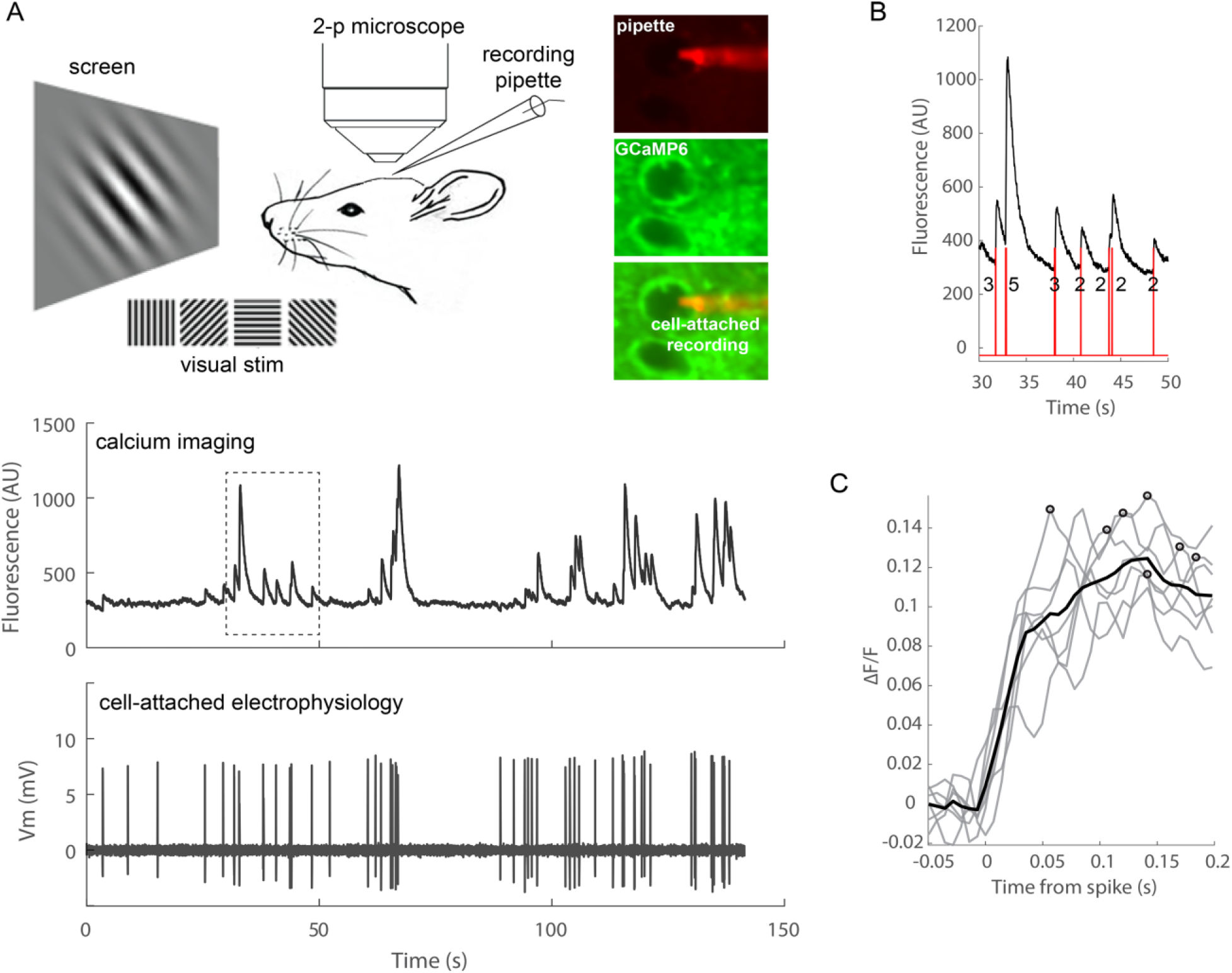
Simultaneous calcium imaging and electrophysiology *in vivo.* (**A**) Experimental design and example fluorescence and Vm traces from an example Emx1-s neuron. (**B**) Spike-triggered fluorescence responses. Red lines indicate spikes and numbers indicate the number of spikes in each spiking event. Trace corresponds to the boxed region in (**A**). (**C**) Fluorescence changes (ΔF/F) in response to representative single-spike events from the recorded cell shown in (**A**). For GCaMP6s, the spike summation window and calcium response window were 150 ms and 200 ms, respectively. Black line indicates mean ΔF/F. Circles indicate peak fluorescence changes (ΔF/F peak) within the calcium response window.

**Table 1.** Dataset. Sample size for each mouse line. Spon and Visstim: spontaneous and visually-evoked recordings, respectively. Total: total number of cells in high zoom dataset. Low zoom: cells imaged at both high and low optical zooms. *One Emx1-f cell was imaged at the same low optical zoom but with smaller FOV and at ~60 fps.

| Mouse line | Acronym | GECI | # Mice | # Cells |  |  |  |
| --- | --- | --- | --- | --- | --- | --- | --- |
|  |  |  |  | Spon | Visstim | Total | Low zoom |
| Emx1-IRES-Cre;Camk2a-tTA;Ai94 | Emx1-s | GCaMP6s | 7 | 21 | 11 | 32 | 6 |
| Camk2a-tTA;tetO-GCaMP6s | tetO-s | GCaMP6s | 2 | 6 | 0 | 6 | 0 |
| Emx1-IRES-Cre;Camk2a-tTA;Ai93 | Emx1-f | GCaMP6f | 6 | 9 | 17 | 26 | 7* |
| Cux2-CreERT2;Camk2a-tTA;Ai93 | Cux2-f | GCaMP6f | 7 | 4 | 23 | 27 | 9 |
| Total |  |  | 22 | 40 | 51 | 91 | 22 |

Recording/imaging sessions of 2-4 min each were obtained from these 237 cells (multiple sessions were collected for some cells in cases where the patch remained stable). We selected 91 cells with high-quality recording and imaging conditions from the 4 mouse lines (**Table 1**) for analysis, each with only one recording/imaging session for unbiased sampling (see **Figure 1-figure supplement 1** for distribution of mouse age, cell depth, and firing rate). Cell selection was based on qualitative assessment of both imaging and electrophysiology data by experienced annotators. Specifically, selected cells exhibited (1) no apparent motion artifacts (after motion correction in x-y axes); (2) no apparent photobleaching; (3) no apparent damage (e.g. becoming filled with dye from the pipet); and (4) stable baseline and distinguishable spikes for electrophysiological recordings.

### Constructing single-cell spike-to-calcium fluorescence response curves

The firing rates of individual cells, computed from the total number of spikes detected during a recording, were comparable across mouse lines (**Figure 1-figure supplement 1C**). We focused on isolated spiking events with calcium responses well-separated from those of adjacent events. A spiking event is defined as a group of spikes within a spike summation window (150 ms and 50 ms for GCaMP6s and GCaMP6f, respectively) with no spikes in the pre-event and post-event exclusion windows (150 ms and 50 ms pre and post respectively for GCaMP6s, and 50 ms both pre and post (after the last spike) for GCaMP6f). We summed the number of spikes within each spiking event, aligned fluorescence responses to the first spike within the event, and computed the peak fluorescence change (ΔF/F peak) during the calcium response window (200 ms for GCaMP6s, 50 ms or 75 ms for GCaMP6f in single-spike or multispike events) (**Figure 1B, C**; see Methods for spike exclusion, spike summation, and calcium response windows). Spiking events contained 1-5 spikes. Events with >5 spikes were excluded from analysis due to the low frequency of such events (≥3 6-spike events were observed in 2 out of 32 Emx1-s cells, 0 out of 6 tetO-s cells, 5 out of 26 Emx1-f cells, and 1 out of 27 Cux2-f cells). 70-78% of detected spikes were in isolated spiking events with ≤5 spikes. >50% of analyzed spikes occurred in multi-spike events and not as isolated single spikes. We constructed single-cell spike-to-calcium fluorescence response curves (ΔF/F peak as a function of the number of spikes, with a minimum requirement of 3 events per bin; **Figure 2**).

**Figure 2.**
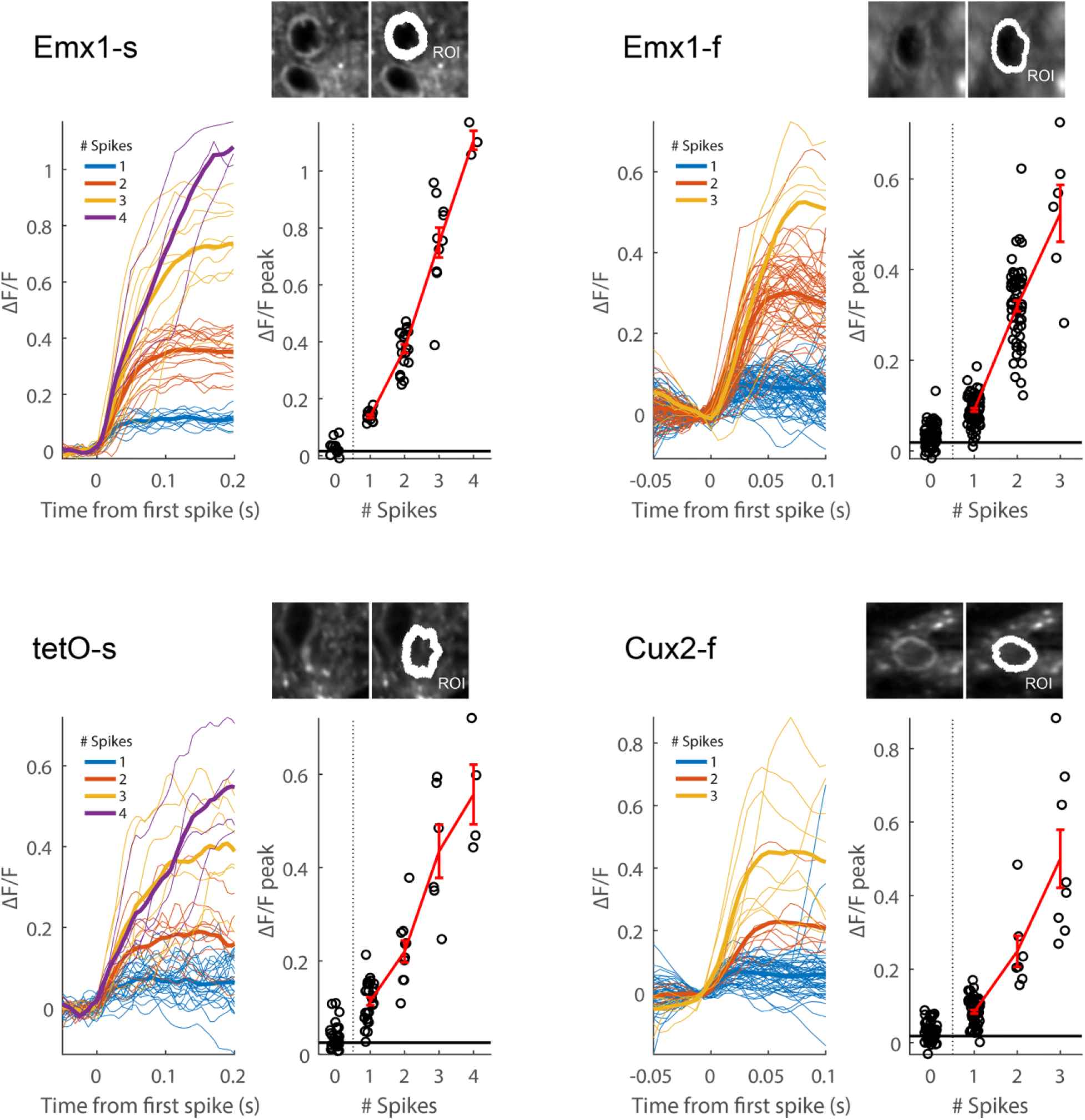
Example single-cell spike-to-calcium fluorescence response curves. For each mouse line: Cell ROI and segmentation mask (top right), ΔF/F traces (bottom left), and spike-to-calcium fluorescence response curve (ΔF/F peak as a function of the number of spikes, with a minimum requirement of 3 events per bin; bottom right). Horizontal black lines indicate the standard deviation of ΔF/F during no-spike intervals. Estimated zero-spike ΔF/F peak were plotted for comparison. Error bars show sem. Spike summation windows were 150 ms and 50 ms for GCaMP6s and GCaMP6f, respectively.

For calcium imaging data, we first examined the effect of fluorescence background subtraction on the peak ΔF/F signal. Following methods for virally expressed GCaMP6 (Chen et al., 2013), the background neuropil signal was estimated as the mean fluorescence of all pixels within 20-μm from the cell center, excluding the selected cell, and neuropil subtraction was performed as: F_corrected_(t) = F_measured_(t) – r × F_neuropil_(t), where r was the neuropil contamination ratio (**Figure 2-figure supplement 1**). Neuropil subtraction using a constant r-value for all cells (ranging from 0.1 to 0.7, where 0.7 was used for virally expressed GCaMP6) did not consistently decrease within-cell variability of ΔF/F peak *(i.e.* variability across spiking events) (**Figure 2-figure supplement 1D**). Given the potential for artificially boosting ΔF/F due to lower baseline fluorescence (see example cell in **Figure 2-figure supplement 1B-C**), we chose not to perform neuropil correction in this study (however, see **Figure 3-figure supplement 1** and **Figure 5-figure supplement 2** below).

### Population spike-to-calcium fluorescence response curves

To compare the relationship between spikes and fluorescence, population response curves were constructed by averaging together responses of individual neurons from each mouse line (**Figure 3A, B**). Spontaneous and visually-evoked responses were similar even though they were not recorded from the same cell and/or animal, and were therefore pooled in the population responses. Data from mice anesthetized with isoflurane (n = 77 cells from 18 mice) and urethane (n = 14 cells from 4 mice) were also pooled as their response curves and firing rates were similar. As expected, neurons in GCaMP6s mice exhibited larger single-spike fluorescence responses (ΔF/F peak: 7.8 ± 2.6% in Emx1-s and 13.6 ± 3.7% in tetO-s, mean ± sd) than neurons containing GCaMP6f (4.5 ± 1.6% in Emx1-f and 6.0 ± 1.6% in Cux2-f) (**Figure 3C, D**). Mean ΔF/F peak in response to 5-spike events was 62% for Emx1-s, 73% for tetO-s, 62% for Emx1-f, and 52% for Cux2-f (**Figure 3B**, bottom).

**Figure 3.**
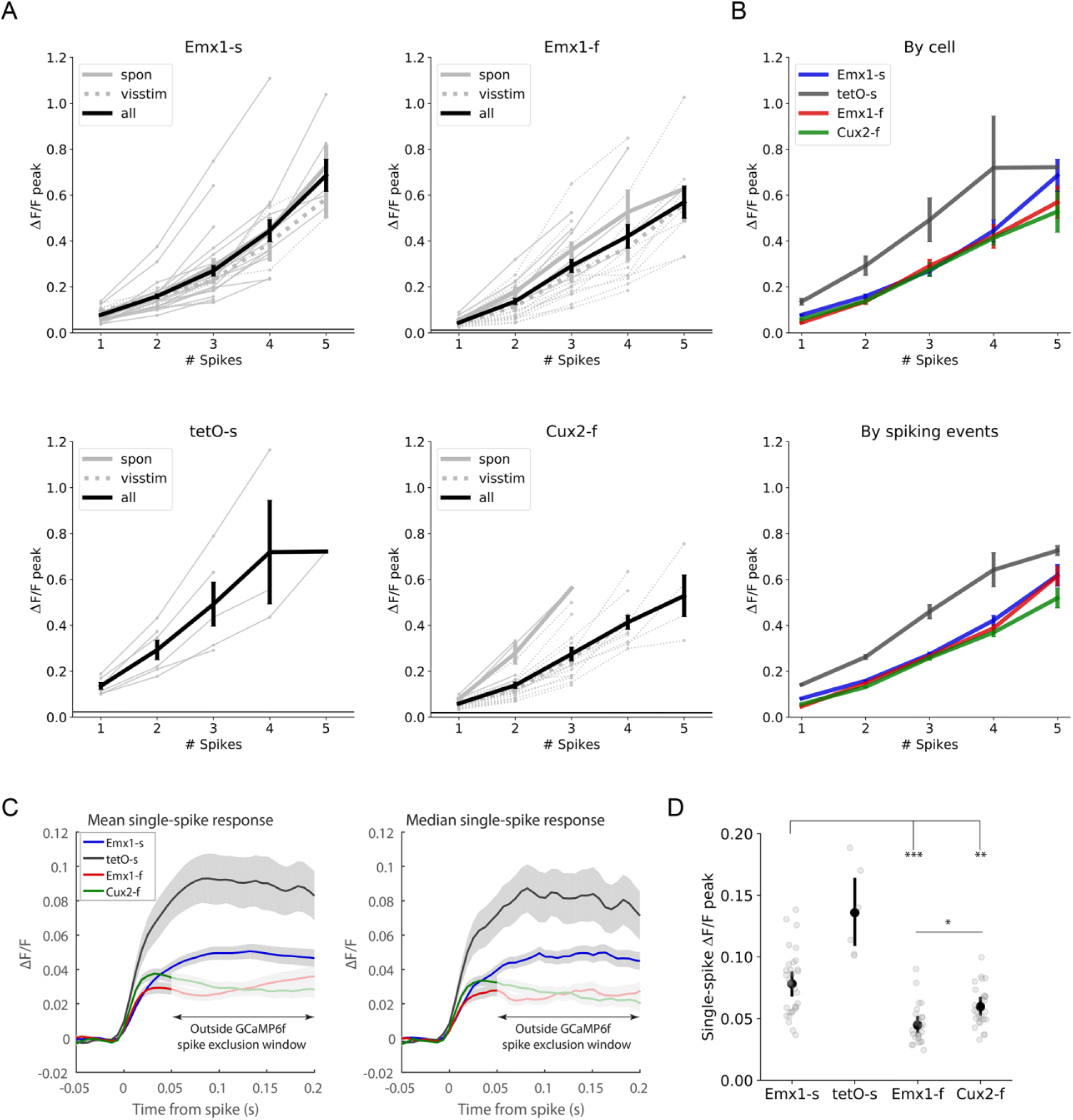
Population response. (**A**) Population spike-to-calcium fluorescence response curves. Each line represents one cell. Bold lines indicate mean responses (± sem) for spontaneous, visually-evoked, and all recordings. Horizontal black lines indicate the mean standard deviation of ΔF/F during no-spike intervals. (**B**) Overlay of population response curves. By cell (top): population means from (A), computed from the mean response of individual cells. Emx1-s: n = 32 cells; tetO-s: n = 6; Emx1-f: n = 26; Cux2-f: n = 27. By spiking events (bottom): population response computed from spiking events pooled from all cells as in Chen et al., 2013. Emx1-s: n = 1160, 599, 255, 103, and 52 spiking events for 1-5 spikes; tetO-s: n = 250, 103, 48, 17, and 6; Emx1-f: n = 1204, 921, 416, 157, and 60; Cux2-f: n = 2667, 809, 209, 118, and 46. (**C**) Mean and median ΔF/F in response to single-spike events. For GCaMP6f, the single-spike calcium response window was 50 ms. Shading corresponds to 2×sem. (**D**) Mean single-spike ΔF/F peak of each cell (p = 1e-7, ANOVA; *, p < 0.05; **, p < 0.01; ***, p ≤ 0.001, multiple comparison test using Tukey’s honest significant difference criterion; error bars show 95% confidence interval around the mean).

For a more direct comparison with that of viral GCaMP6 (Chen et al., 2013), we analyzed our data using a neuropil contamination ratio of 0.7 (**Figure 3-figure supplement 1**). Aggregating spiking events across cells, mean ΔF/F peak in response to 5-spike events within 150 ms and 50 ms for GCaMP6s and GCaMp6f, respectively, was 135% for Emx1-s, 117% for tetO-s (130% in response to 4-spike events), 130% for Emx1-f, and 93% for Cux2-f. In comparison, Chen et al. (2013) reported ~200% and ~100% mean ΔF/F peak in response to 5-spike events within 250 ms for viral GCaMP6s and GCaMP6f, respectively.

### Fluorescence response variability

GCaMP6f-containing neurons exhibited more within-cell variability of ΔF/F peak than GCaMP6s-containing neurons. The coefficient of variation across spiking events, a measure of within-cell variability, was greater in Emx1-f and Cux2-f than Emx1-s and tetO-s (**Figure 4A**). Single-spike events exhibited larger within-cell variability than multi-spike events only in GCaMP6f mice (**Figure 4B**). Between-cell (cell-to-cell) variability of ΔF/F peak was generally similar across the 4 mouse lines (**Figure 4C**).

**Figure 4.**
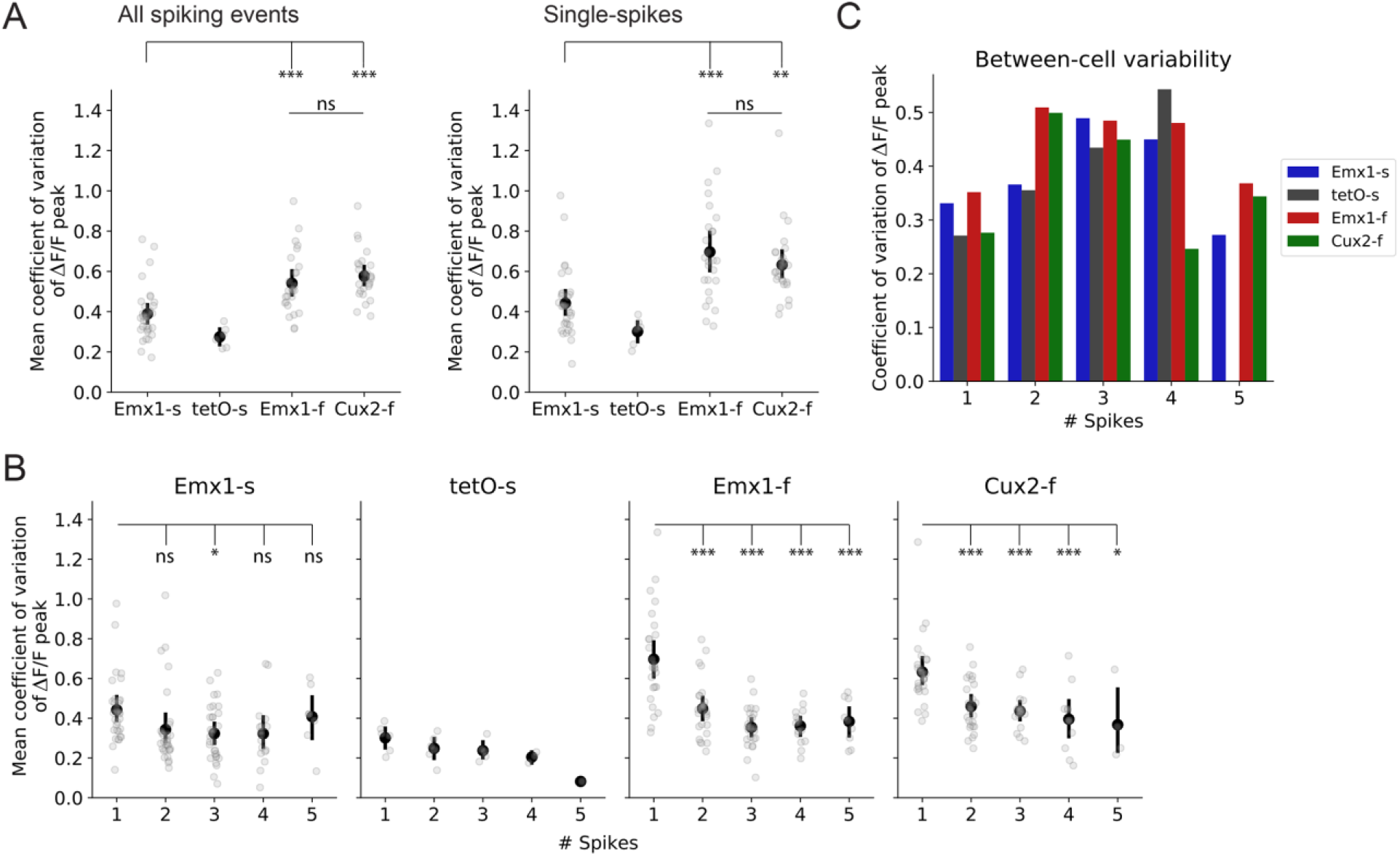
Within-cell and between-cell variability of ΔF/F peak. (**A**) Within-cell variability. Mean coefficient of variation of ΔF/F peaks of each cell computed from all spiking events (left panel; p = 1e-6, ANOVA) or single-spike events (right panel; p = 9e-6, ANOVA). (**B**) Within-cell variability (coefficient of variation of ΔF/F peak) for each spike bin (Emx1-s: p = 0.04; tetO-s: p = 0.03; Emx1-f: p = 3e-11; Cux2-f: p = 6e-6, ANOVA). ns, p > 0.05; *, p < 0.05; **, p < 0.01; ***, p ≤ 0.001, multiple comparison test using Tukey’s honest significant difference criterion. Error bars show 95% confidence interval around the mean. (**C**) Cell-to-cell variability. Coefficient of variation of ΔF/F peak from the population response in Fig. 3B (top). Spike summation windows were 150 ms and 50 ms for GCaMP6s and GCaMP6f, respectively.

We found that as expected, photon shot noise was the dominant noise source in images from all mouse lines. The slope of the least squares fit between the variance and mean of the number of photons over time for all pixels in the FOV was 1.03 ± 0.04 (mean ± sd), consistent with the noise following a Poisson process (with intercept of −0.06 ± 0.13, 91 cells; example cell in **Figure 4-figure supplement 1A**). Furthermore, most of the trial-to-trial variance in the amplitudes of calcium transients was attributable to shot noise, with trial-to-trial variance being only fractionally greater than expected from shot noise (measured variance greater than variance expected from shot noise by 1 ± 14% in Emx1-s, 0 ± 8% in tetO-s, 2 ± 7% in Cux2-f; mean ± sd, **Figure 4-figure supplement 1B**). The exception was Emx1-f, in which the trial-to-trial variance was ~50% greater than expected from shot noise (48 ± 91%).

The signal-to-noise ratio (SNR) was greater in GCaMP6s than GCaMP6f lines. We determined the noise floor in no-spike intervals of ≥1 s, separated by ≥4 s and ≥1 s from previous and subsequent spikes, respectively, and sampled the corresponding calcium fluorescence traces randomly to obtain snippets of fluorescence with the same length as single-spike events, computing the standard deviation of ΔF/F as an estimate of noise. The single-spike SNR (peak ΔF/F divided by standard deviation of no-spike ΔF/F) for Emx1-s was greater than for Emx1-f and Cux2-f (**Figure 4-figure supplement 2**).

### Spike detection when imaging a single neuron, at high zoom

GCaMP6 indicators have been widely adopted because they exhibit greater spike-evoked ΔF/F than previous GCaMP indicators, but still some spikes may go undetected (Chen et al., 2013). We found that, on average, the majority of 2-spike events were detected in all four mouse lines and the majority of 1-spike events in GCaMP6s lines. Mean (± sd) spike detection rates for 1- and 2-spike events were 70 ± 20% and 96 ± 6% for Emx1-s, 91 ± 11% and 100% for tetO-s, 49 ± 18% and 88 ± 18% for Emx1-f, and 43 ± 23% and 76 ± 18% for Cux2-f, at 5% false positive rate (**Figure 5**). There were substantial cell-to-cell differences in spike detection rates in both GCaMP6s and GCaMP6f lines (single spike detection rate ranges 31-100% and 11-93% for GCaMP6s and GCaMP6f, at 5% false positive rate, **Figure 5**). Single-spike detection rates were high in some GCaMP6s neurons, but few GCaMP6f neurons (≥90% detection at 5% false positive rate in 20% (6 of 30) Emx1-s cells, 75% (3 of 4) tetO-s cells, 4% (1 of 23) Emx1-f cells and 0% (0 of 23) Cux2-f cells).

**Figure 5.**
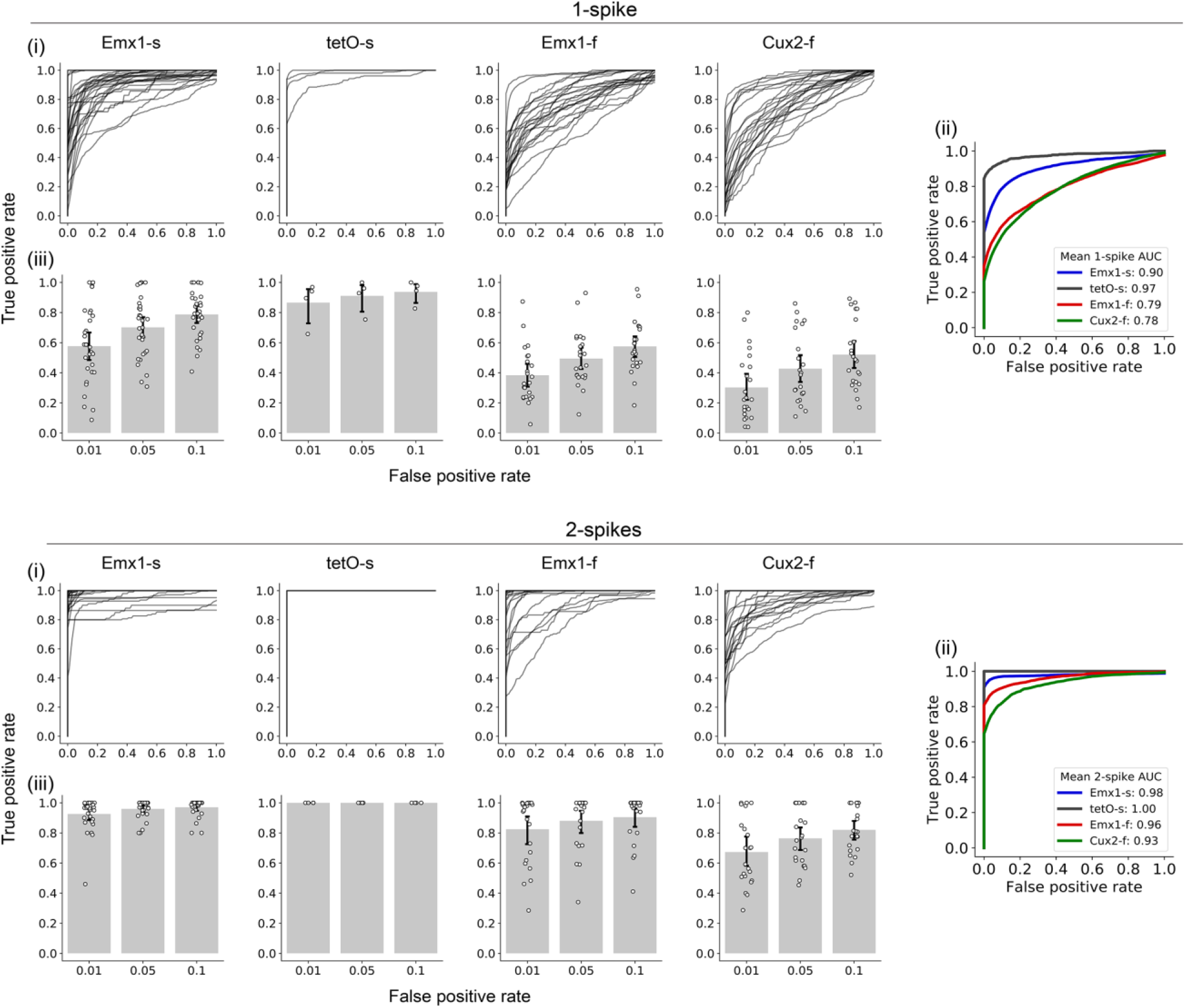
Spike detection when imaging a single neuron, at high zoom. (i) Receiver operating characteristic (ROC) curves for classifying 1-spike (top) and 2-spike (bottom) events in individual cells. (ii) Mean detection rate across cells. (iii) Detection rate (true positive rate) at 1%, 5%, and 10% false positive rate. Error bars show 95% confidence interval around the mean (Emx1-s: n = 30 cells with ≥1 no-spike interval out of 32 total cells; tetO-s: n = 4 out of 6; Emx1-f: n = 23 out of 26; Cux2-f: n = 23 out of 27).

Previous studies have documented substantial contamination of somatic traces with fluorescence from the surrounding neuropil, due to the extended nature of the microscope point spread function. Neuropil contamination is often removed by subtracting a scaled version of the neuropil fluorescence from the somatic fluorescence, with the scale factor often referred to as the r-value (Akerboom et al., 2012). Some studies have employed the same r-value for all neurons, while others, including the Allen Brain Observatory (de Vries et al., 2019), have tuned the r-value for each neuron, reasoning that the fluorescence of the neuropil varies across space, near blood vessels for example.

For each neuron, we found the optimal r-value (resulting in the maximum number of detected spikes at a fixed false positive rate, **Figure 5-figure supplement 1**). Optimal r-values differed greatly between cells. Mean (± sd) optimal r-values were 0.60 ± 0.39 for Emx1-s, 0.50 ± 0.27 for tetO-s, 0.47 ± 0.38 for Emx1-f, and 0.42 ± 0.44 for Cux2-f (**Figure 5-figure supplement 2**). These r-value ranges are similar to those employed in the Allen Brain Observatory for these mouse lines. At optimal r-values, single-spike detection rates (77 ± 18% for Emx1-s, 93 ± 10% for tetO-s, 51 ± 19% for Emx1-f, and 43 ± 23% for Cux2-f, at 5% false positive rate) were slightly greater than in the absence of neuropil subtraction (**Figure 5-figure supplement 2B**). Hence neuropil contamination varies substantially across cells in all four mouse lines and tuning the neuropil subtraction for each neuron slightly improves single spike detection.

### Spike detection during population imaging, at low zoom

Many population calcium imaging experiments are performed at low optical zoom, offering a large field of view containing many neurons. The laser dwells for a shorter time on each neuron at low zoom than at high zoom, generating fewer fluorescence photons, resulting in lower SNR and a reduced spike detection rate.

For a subset of neurons, we acquired images at high and low zoom, the latter yielding ~400 x 400 μm field of view, comparable to that commonly employed in population-scale studies such as the Allen Brain Observatory (**Table 1, Figure 6A, B**). We compared numbers of emitted photons and spike detection at high and low zoom, using identical laser powers to facilitate comparison.

**Figure 6.**
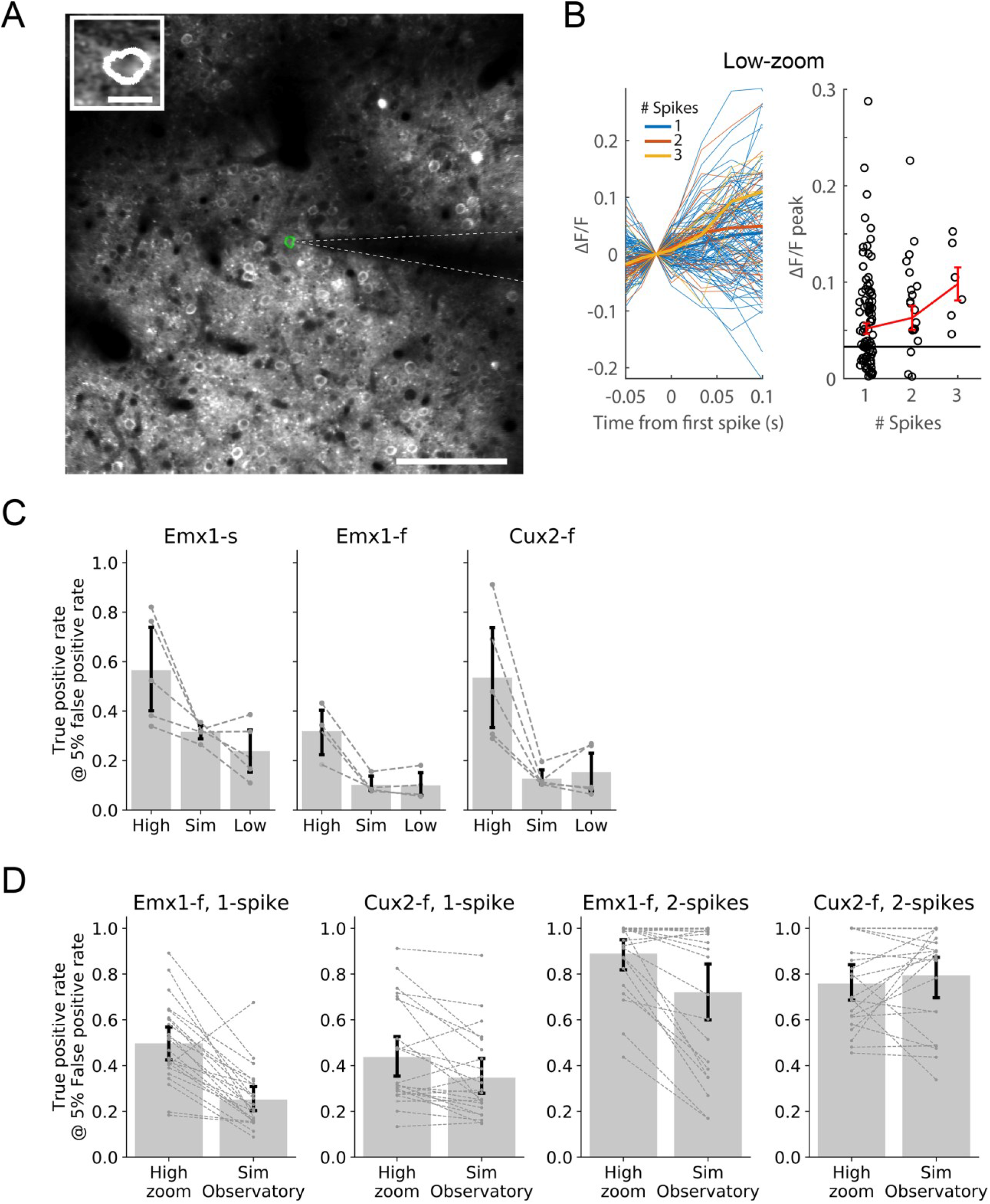
Spike detection during population imaging, at low zoom. (**A**) FOVs and cell ROIs for an example Cux2-f neuron imaged at both high (white ROI, inset) and low (green ROI) optical zooms. Scale bars, high zoom: 10 μm; low zoom: 100 μm. (**B**) ΔF/F traces and spike-to-calcium fluorescence response curves at low zoom for the neuron shown in (A). Horizontal black line indicates the standard deviation of ΔF/F during no-spike intervals. Error bars show sem. (**C**) Spike detection using measured high zoom traces (High), simulated low zoom traces (Sim) and measured low zoom traces (Low). Error bars show 95% confidence interval around the mean (Emx1-s: n = 5 cells with ≥1 no-spike interval for both high and low zoom; Emx1-f, n = 4; Cux2-f, n = 5). (**D**) Expected spike detection for Emx1-f and Cux2-f lines in the Allen Brain Observatory. Error bars show 95% confidence interval around the mean. Brain Observatory was simulated using 22 high zoom Emx1-f cells and 23 Cux2-f cells with ≥1 no-spike interval.

The expected change in photon flux, upon changing the zoom, is the product of the change in field of view (ratio of areas = 455) and the change in pixel dwell time. As in many commercial multiphoton instruments, the dwell time was set automatically by our 2-photon microscope. With a homogenous fluorescent slide, we measured a high/low photon flux ratio of 323 ± 19, corresponding to a dwell time ratio of 0.71 ± 0.04. The measured photon flux from calcium imaging data was 5.03 x 10^5^ ± 3.72 x 10^5^ photons per cell per second at high zoom and 2,785 ± 2,216 at low zoom (see Methods), corresponding to a high/low ratio of 333 ± 430 (22 cells with high and low zoom data). Hence the change in photon flux from cortex, when changing the microscope field of view, was as expected.

Spike detection was compromised at low zoom, by a factor expected from the decrease in photon flux. Measured single spike detection rates were 57 ± 22% at high zoom and 24 ± 11% at low zoom in Emx1-s, 32 ± 10% and 10 ± 6% in Emx1-f, and 54 ± 27% and 15 ± 10% in Cux2-f (at 5% false positive rate, **Figure 6C**). We calculated expected detection rates at low zoom by adding noise to our high zoom traces, to mimic the change in photon flux and resulting shot noise upon switching from high to low zoom (see Methods). Expected spike detection rates at low zoom were 32 ± 3% in Emx1-s, 10 ± 4% in Emx1-f, and 13 ± 4% in Cux2-f, not significantly different from measured rates for any of the three mouse lines (p > 0.05, paired sample t-test, **Figure 6C**).

Maintaining laser power when switching from high to low zoom in our study resulted in lower laser power and fewer fluorescent photons at low zoom than in typical population imaging studies and we expect spike detection rates during typical population imaging to be between those measured in our high and low zoom experiments. In the Allen Brain Observatory, photon flux was 30,341 ± 14,816 photons per neuron per second in Emx1-f (4,160 neurons) and 31,779 ± 20,526 in Cux2-f (7,441 neurons). We calculated the expected spike detection rates for the Allen Brain Observatory, at 5% false positive rate, to be 25 ± 13% for Emx1-f and 35 ± 19% for Cux2-f for single spikes and 72 ± 32% for Emx1-f and 79 ± 22% for Cux2-f for 2-spike events (**Figure 6D**). Thus, most spiking events with ≥2 spikes are likely to be detected. Pixel dwell time, frame rate and laser power used in the Allen Brain Observatory are comparable to those commonly used in many 2-photon experiments and we expect these calculated single- and two-spike detection rates are likely reasonable estimates of spike detection with GCaMP6 indicators in many population imaging studies.

## Discussion

Calcium imaging is widely used to report neuronal spiking activity *in vivo*. However, accurate spike inference from calcium imaging remains a challenge, and there are relatively few ground truth datasets with simultaneous calcium imaging and electrophysiology to aid the development of more accurate spike inference algorithms. In a recent challenge (Spike Finder; http://spikefinder.codeneuro.org/) (Berens et al., 2018), ~40 algorithms were trained and tested on datasets consisting of 37 GCaMP6-expressing cells, underscoring the need for additional GCaMP6 calibration data. In addition to supporting efforts toward spike inference, an improved understanding of the relationship between spiking and observed fluorescence signal is necessary to further broaden the utility and impact of calcium imaging. To these ends, we contribute a ground truth dataset consisting of 91 V1 L2/3 excitatory neurons recorded at singlecell resolution (available at https://portal.brain-map.org/explore/circuits/oephys), and characterized their spike-to-calcium fluorescence transfer function. Complementing existing datasets with viral GECI expression (Chen et al., 2013; Theis et al., 2016; Dana et al., 2016), our work facilitates interpretation of existing and future calcium imaging studies using mainstream transgenic mouse lines, such as the Allen Institute’s Brain Observatory Visual Coding dataset (http://observatory.brain-map.org/visualcoding) (de Vries et al., 2019).

We found that shot noise was the dominant noise source in our imaging experiments for most cells. Furthermore, most of the trial-to-trial variance in the amplitudes of calcium transients was attributable to shot noise, with trial-to-trial variance being on average only ~2% greater than expected from shot noise in Emx1-s, tetO-s, and Cux2-f. This additional variance is presumably the cumulative effects of motion, instrumentation noise and spike-to-calcium coupling. We conclude that there’s little trial-to-trial variability in spike-to-calcium coupling in 3 of our 4 mouse lines. The exception was Emx1-f, in which the trial-to-trial variance was ~50% greater than expected from shot noise. We suspect the greater variability in Emx1-f may be biological and possibly related to this line’s susceptibility to epileptiform activity (Steinmetz et al., 2017), and we explore this topic further in our follow-up paper (Ledochowitsch et al., https://www.biorxiv.org/content/10.1101/800102v1).

Our results report reliable single spike detection in transgenic mouse lines expressing GCaMP6s, when imaged at high zoom. Single spike detection rates were variable across cells but were ~70% on average and ≥90% in some neurons, at 5% false positive rate. Effective single spike detection is consistent with results from virally-expressed GCaMP6s imaged at high zoom (Chen et al., 2013). Single spike detection rates were generally lower in GCaMP6f expressing mice (~40-50% at 5% false positive rate).

We tested if spike detection rate was dependent on the neuropil contamination ratio r. We found optimal r-values for maximizing single-spike detection rates in individual cells. The effect of optimizing r-value was most pronounced in Emx1-s, where optimal neuropil subtraction increased the single spike detection rate by ~7% at 5% false positive rate. The increased detection rate is a potential benefit of tuning r-values for individual neurons, as performed for our Brain Observatory dataset (de Vries et al., 2019), over using the same r-value for all neurons.

As expected, single spike detection rates were far lower at low zoom (~10-20% at 5% false positive rate), where the neuron occupies only a small percentage of the field of view, than at high zoom where the cell almost fills the field of view. Our low zoom images, with a 400 x 400 μm field of view imaged at 30 fps, are similar to many population imaging experiments, suggesting that spike detection rates in many population calcium imaging studies may be closer to 10% than 70%. That said, our low zoom images had lower median photon flux than the Allen Brain Observatory, and perhaps of many typical population imaging experiments, a result of lower illuminating laser power used. Hence ~10-20% single spike detection rate may be lower than in many typical population imaging experiments. Indeed, when we adjusted our high zoom data to mimic the typical photon flux level found in the Allen Brain Observatory data, we found the estimated single spike detection rates to be 25-35% for GCaMP6f cells.

In summary, in this study we present a ground truth dataset with simultaneous electrophysiology and calcium imaging. We compared multiple aspects of spike-response properties between different GECIs (GCaMP6s and GCaMP6f) and among several transgenic lines. For example, we found that GCaMP6s and GCaMP6f cells have similar spike-fluorescence response curves, but GCaMP6f cells have greater variability and lower single-spike detection rates than GCaMP6s cells. Additionally, this dataset is well suited for testing novel methods, including but not limited to data resampling (e.g. to approximate the spatiotemporal resolution and noise profile of population-scale imaging experiments), data preprocessing (e.g. neuropil subtraction) and spike inference. By making our data freely available, we hope that it will serve the community as a further resource to better understand quantitatively the link between calcium-evoked fluorescent imaging signals and spiking activity.

## Materials and Methods

### Key Resources Table

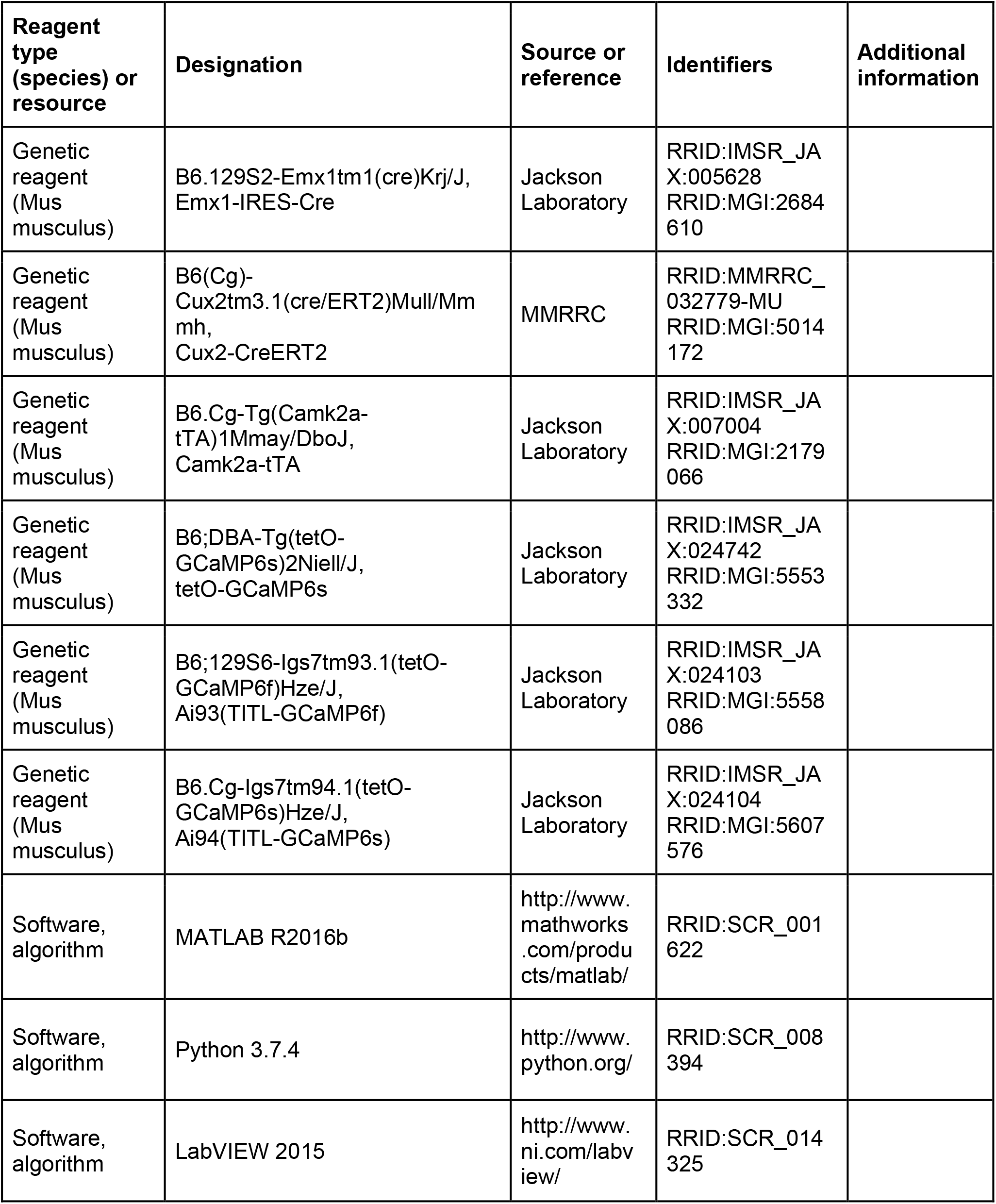

Experimental procedures were in accordance with NIH guidelines and approved by the Institutional Animal Care and Use Committee (IACUC) of the Allen Institute for Brain Science. All experiments were conducted at the Allen Institute for Brain Science.

#### Mice

2-p-targeted electrophysiology and 2-p calcium imaging was conducted in adult transgenic mice (2-6 months old, both sexes, n = 22), including *Emx1*-IRES-Cre;*Camk2a*-tTA;Ai94 (simplified as Emx1-s, n = 7), *Camk2a*-tTA;tetO-GCaMP6s (Wekselblatt et al., 2016) (simplified as tetO-s, n = 2), *Emx1*-IRES-Cre;*Camk2a*-tTA;Ai93 (simplified as Emx1-f, n = 6), and *Cux2*-CreERT2;*Camk2a*-tTA;Ai93 (simplified as Cux2-f, n = 7). Ai93 and Ai94 containing mice (Madisen et al., 2015) included in this dataset did not show behavioral signs for epileptic brain activity (Steinmetz et al., 2017).

#### Surgery

Mice were anesthetized with either isoflurane (0.75-1.5% in O_2_) or urethane (1.5 g/kg, 30% aqueous solution, intraperitoneal injection), then implanted with a metal head-post. A circular craniotomy was performed with skull thinning over the left V1 centering on 1.3 mm anterior and 2.6 mm lateral to the Lambda. During surgery, the craniotomy was filled with Artificial Cerebrospinal Fluid (ACSF) containing (in mM): NaCl 126, KCl 2.5, NaH_2_PO_4_ 1.25, MgCl_2_ 1, NaHCO_3_ 26, glucose 10, CaCl_2_ 2, in ddH_2_O; 290 mOsm; pH was adjusted to 7.3 with NaOH to keep the exposed V1 region from overheating or drying. Durotomy was performed to expose V1 regions of interest that were free of major blood vessels to facilitate the penetration of recording micropipettes. A thin layer of low melting-point agarose (1-1.3% in ACSF, Sigma-Aldrich) was then applied to the craniotomy to control brain motion. The mouse body temperature was maintained at 37°C with a feedback-controlled animal heating pad (Harvard Apparatus).

#### Calcium imaging

Individual GCaMP6+ neurons within ~100-300 μm underneath the pial surface were visualized under adequate depth of anesthesia (Stage III-3) using a Bruker (Prairie) 2-p microscope with 8 kHz resonant-galvo scanners, coupled with a Chameleon Ultra II Ti:sapphire laser system (Coherent). Fluorescence excited at 920 nm wavelength, with <70 mW laser power measured after the objective, was collected in two spectral channels using green (510/42 nm) and red (641/75 nm) emission filters (Semrock) to visualize GCaMP6 and the Alexa Fluor 594-containing micropipette, respectively. Fluorescence images were acquired at various framerates (~118-158 fps, and additionally at ~30 or 60 fps for subset of cells) through a 16x water-immersion objective lens (Nikon, NA 0.8), with or without visual stimulations.

#### Electrophysiology

2-p targeted cell-attached recording was performed following established protocols (Margrie et al., 2003; Kitamura et al., 2008; Knoblich et al., 2019). Long-shank borosilicate (KG-33, King Precision Glass) micropipettes (5-10 MΩ) were pulled with a P-97 puller (Sutter) and filled with ACSF and Alexa Fluor 594 to perform cell-attached recordings on GCaMP6+ neurons. Micropipettes were installed on a MultiClamp 700B headstage (Molecular Devices), which was mounted onto a Patchstar micromanipulator (Scientifica) with an approaching angle of 31 degrees from horizontal plane. Minimal seal resistance was 20 MΩ. Data were acquired under “I = 0” mode (zero current injection) with a Multiclamp 700B, recorded at 40 kHz using Multifunction I/O Devices (National Instruments) and custom software written in LabVIEW (National Instruments) and MATLAB (MathWorks). Isoflurane level was intentionally adjusted during recording sessions to keep the anesthesia depth as light as possible, resulting in fluctuation of the firing rates of recorded neurons.

#### Visual stimulation

Whole-screen sinusoidal static and drifting gratings were presented on a calibrated LCD monitor spanning 60° in elevation and 130° in azimuth to the contralateral eye. The mouse’s eye was positioned ~22 cm away from the center of the monitor. For static gratings, the stimulus consisted of 4 orientations (45° increment), 4 spatial frequencies (0.02, 0.04, 0.08, and 0.16 cycles per degree), and 4 phases (0, 0.25, 0.5, 0.75) at 80% contrast in a random sequence with 10 repetitions. Each static grating was presented for 0.25 seconds, with no inter-stimulus interval. A gray screen at mean illuminance was presented randomly a total of 60 times. For drifting gratings, the stimulus consisted of 8 orientations (45° increment), 4 spatial frequency (0.02, 0.04, 0.08, and 0.16 cycle per degree) and 1 temporal frequency (2 Hz), at 80% contrast in a random sequence with up to 5 repetitions. Each drifting grating lasted for 2 seconds with an inter-stimulus interval of 2 seconds. A gray screen at mean illuminance was presented randomly for up to 15 times.

#### Data analysis

Electrophysiology and calcium imaging data were analyzed using custom MATLAB and Python scripts. For electrophysiology, Vm was filtered between 250 Hz and 5 kHz, and automated spike detection was performed using a threshold criterion (5×std of Vm). Unusually prolonged transient increases in calcium fluorescence were excluded from analysis with an adaptive Vm threshold: 0.25×V_baseline_+(V_peak_-V_baseline_), where V_baseline_ was the mean Vm over 2 ms before the first spike of the spiking event and V_peak_ was the amplitude of the first spike of the spiking event. The cumulative time above this Vm threshold was compared against a time threshold (6 ms × the number of spikes within the spiking event), and the fluorescence associated with spiking events with cumulative time above the time threshold were not analyzed.

For calcium imaging, in-plane motion artifacts were corrected (Dombeck et al., 2007), and cell/ ROI selection was performed using a semi-automatic algorithm (Chen et al., 2013) (kindly provided by Karel Svoboda, Janelia Research Campus). Ring-shaped ROIs were used to select GCaMP6-positive excitatory neurons, with GCaMP6 expression typically excluded from the nucleus and restricted to the cytoplasm. For 2 of 91 cells, a satisfactory ring-shaped ROI could not be found automatically, and a circular ROI covering the entire soma was used instead. Calcium imaging data was smoothed using a local regression method using weighted linear least squares and a 1^st^ degree polynomial model (built-in “smooth” function in MATLAB with “rlowess” method), and an averaging window of 5 frames.

To construct spike-calcium fluorescence response curves, we first identified all isolated spiking events. For GCaMP6s, isolated spiking events were separated from previous and subsequent spiking events by >150 ms and >50 ms, respectively. For GCaMP6f, isolated spiking events were separated from previous and subsequent spiking events by >50 ms. Within each spiking event, spike summation windows were 150 ms and 50 ms for GCaMP6s and GCaMP6f, respectively, chosen based on the rise time of the GECIs (Chen et al., 2013). To determine spike-triggered calcium fluorescence responses, fluorescence traces were aligned to the first spike in each spiking event. The change in fluorescence, ΔF/F, for each spiking event was calculated as (F-F_0_)/F_0_, where F_0_ was computed locally as the mean fluorescence over 50 ms and 20 ms before the first spike for GCaMP6s and GCaMP6f, respectively. For GCaMP6s, peak ΔF/F was found within 200 ms after the first spike. For GCaMP6f, peak ΔF/F was found within 50 ms and 75 ms after the first spike for single-spike and multi-spike events, respectively. Bursts of >5 spikes were excluded from analysis due to the low frequency of such events. Alignment jitter intrinsic to the imaging frame rate was ≤6.3 ms in 86 of 91 cells (imaged at 158 fps), with mean expected error of 3.2 ms, and between ≤5.6 ms to ≤8.5 ms in others (imaged at 118-179 fps), with mean expected error of 2.8 to 4.3 ms.

To estimate imaging noise, we found no-spike intervals of ≥1 s, separated by ≥4 s and ≥1 s from previous and subsequent spikes, respectively, and randomly sampled the m corresponding calcium fluorescence traces n times, where m×n approximately matched the number of single-spike events, for a minimum of 25 times. This process was repeated 10 times using different random seeds. Noise floor was computed as the standard deviation of the resultant ΔF/F trace. Noise (zero-spike) ΔF/F peak (plotted for comparison in **Figures 2 and 6B**) was computed using the same calcium response windows as single-spike events (50 ms and 200 ms for GCaMP6f and GCaMp6s, respectively).

To quantify the efficiency of detecting spiking events at a single-trial level, we compared fluorescence traces of the response (single-spike events or 2-spike events) to that of imaging noise (estimated as described above). For each cell, the mean response trace was used as the template vector. The template vector was normalized after subtracting the mean to create the unit vector, and the scalar results of projecting the response and noise traces on the unit vector were computed: *r_i_* and *n_i_* for response and noise scalars, respectively. The detection threshold was defined as the x^th^ percentile of *n_i_* values, where 1-x represented the false positive rate (e.g., x = 95 for 95^th^ percentile or 5% false positive rate), and the detection rate (true positive rate) was the fraction of *r*_i_ values above the detection threshold.

To compare the change in photon flux when changing optical zoom, the number of photons was calculated directly from the fluorescence data, taking into account the gain and offset of the data acquisition (see https://github.com/AllenInstitute/QC_2P). Because pixel dwell time in resonant scanning mode may be variable across the x-axis, photon gain and offset were computed pixel-by-pixel along the x-axis (i.e. from image columns instead of the entire FOV). This adjustment allowed for more accurate comparisons of the number of photons within cell ROIs between high zoom and low zoom. (At high zoom, the cell occupied a larger percentage of the FOV and was more affected by the change in photon gain. At low zoom, the cell occupied a small percentage of the FOV, generally close to the center where photon gain was maximum.) We compared the median number of photons of all pixels at high and low zoom in cells and in a homogenous fluorescent slide.

To calculate spike detection rates in simulated low zoom conditions, we first identified 1-spike and 0-spike traces under high zoom conditions, as described above. Fluorescence was converted to number of photons using photon gain and offset calculated from the entire FOV. n traces for each condition were averaged to give the mean number of photons through time for 1- and 0-spike events at high zoom. These two mean traces were scaled by the low/high zoom photon flux ratio (mean number of photons in cell ROI, median over time, in units of photons per cell per second, at low and high zoom). The resulting simulated low zoom mean traces represented the mean number of photons through time for simulated 1- and 0-spike events at low zoom and were on the time scale of high zoom images (~158 fps). Since shot noise is the dominant source of noise, we used Poisson statistics (stdev = sqrt of the mean) to calculate the distribution of noise values for each time point in each trace. To generate a simulated trace, we selected a value for each time point from the time point’s Poisson distribution. We generated 1,000 simulated 1-spike and 1,000 0-spike traces and used these traces to calculate the detection rate and false positive rate, as described above. Detection rates in simulated low zoom conditions were compared against measured low zoom data. We upsampled the measured low zoom traces to the high zoom frame rate (~158 fps) by linear interpolation and scaled the traces such that the integral of the trace remained unchanged by upsampling.

To calculate expected spike detection rates under generally used population imaging conditions, we selected Emx1-f and Cux2-f cells in L2/3 (depth = 175 μm) from the Allen Brain Observatory (http://observatory.brain-map.org/visualcoding). We converted 1-spike and 0-spike fluorescence traces under high zoom conditions to number of photons, using photon gain and offset calculated from the entire FOV. Scaling of mean traces, generation of simulated traces, and spike detection were as described above for calculating spike detection in simulated low zoom conditions. Mean 1-spike and 0-spike traces for each high zoom cell was scaled by the mean Observatory/high zoom photon flux ratio (mean number of photons in cell ROI, median over time, in units of photons per cell per second, for the Observatory and at high zoom).

## Acknowledgements

We are grateful for the Animal Care, Transgenic Colony Management, and Lab Animal Services teams for mouse husbandry, and Carol Thompson and John Phillips for providing project management support. We thank Karel Svoboda, Hod Dana and Tsai-Wen Chen for sharing analysis software. This work was funded by the Allen Institute for Brain Science. This work was also supported by grants from National Natural Science Foundation of China (NSFC31871055) and Guangdong Science and Technology Department (2017B030314026 and 2018B030334001) to L.L. We thank the Allen Institute founders, Paul G. Allen and Jody Allen, for their vision, encouragement, and support.

## Data availability

The dataset is available at https://portal.brain-map.org/explore/circuits/oephys.

---

**Figure 1-figure supplement 1.**
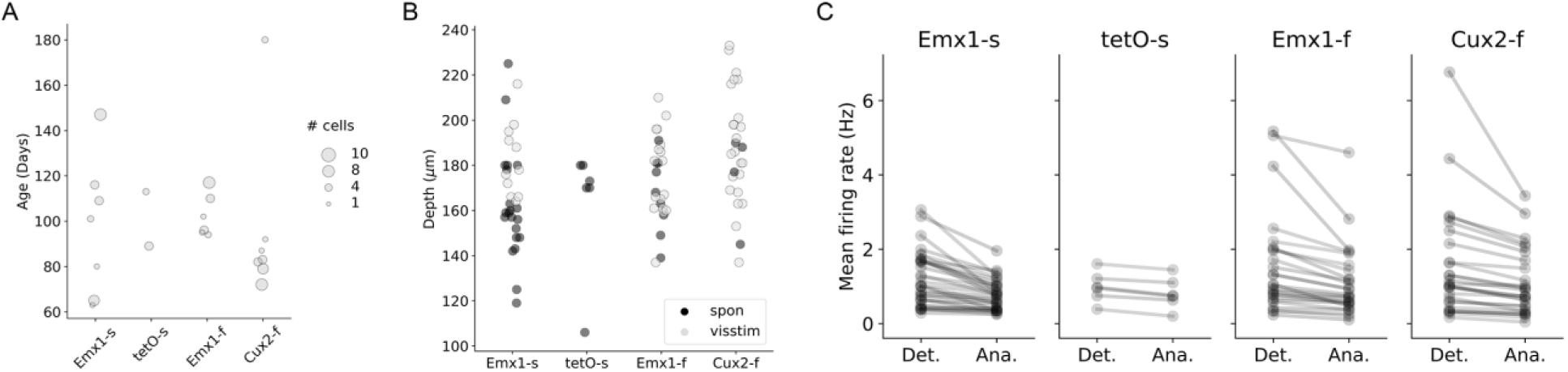
Mouse age, cell depth, and firing rate. (**A**) Mouse age. (**B**) Cell depths. Spon and Visstim: spontaneous and visually-evoked recordings, respectively. (**C**) Mean firing rate computed from all detected spikes (Det.) or analyzed spikes after preprocessing (Ana.; preprocessing included selecting isolated spiking events with ≤5 spikes within spike summation windows of 150 ms and 50 ms for GCaMP6s and GCaMp6f, respectively, and excluding atypical spiking events - see Methods).

**Figure 2-figure supplement 1.**
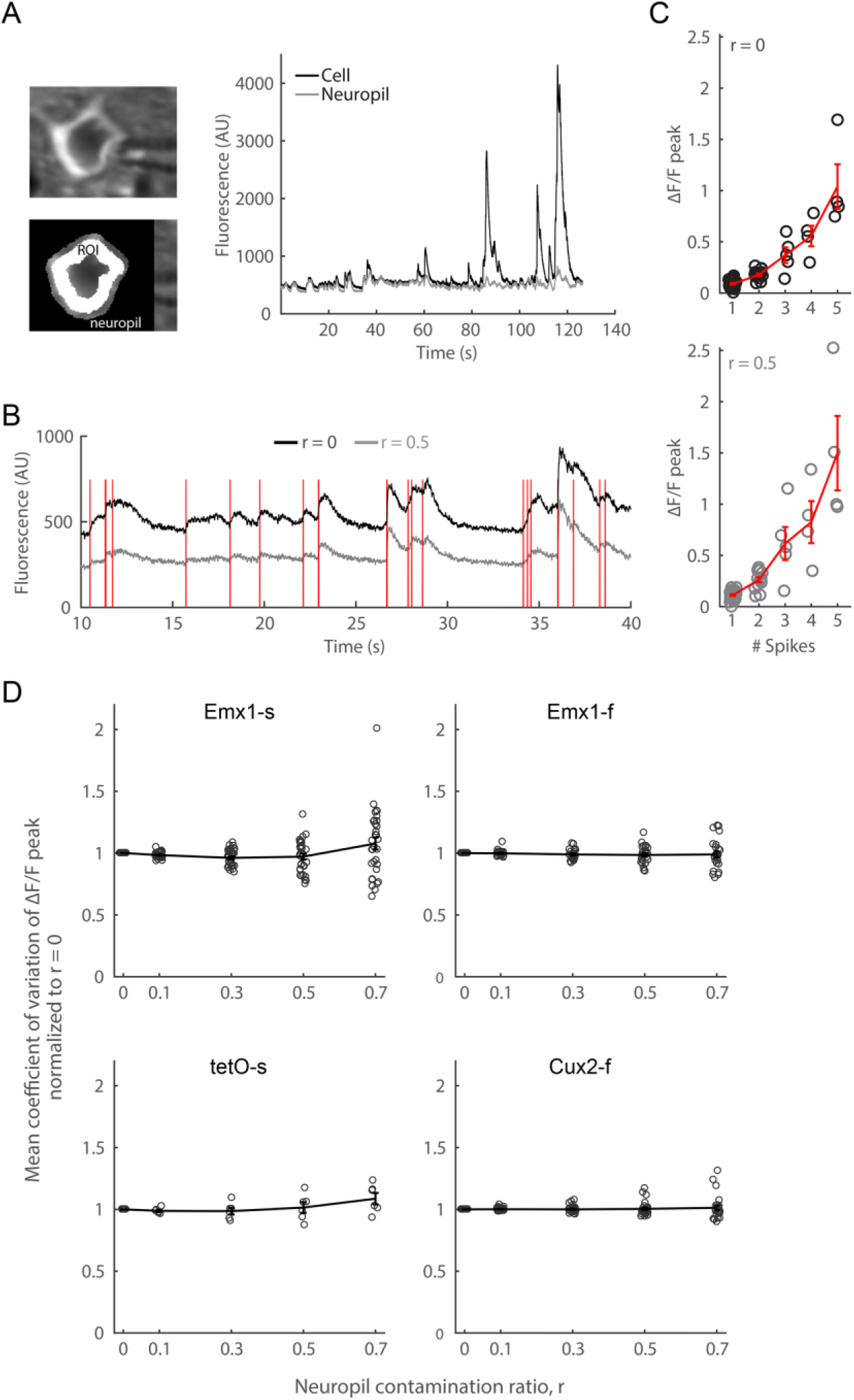
Effect of neuropil subtraction on within-cell variability of ΔF/F peak. (**A**) An example Emx1-s neuron, with labeled regions where cell and neuropil fluorescence were measured, and the corresponding mean fluorescence traces. (**B**) Comparison of cell fluorescence without and with neuropil correction, where the contamination ratio r = 0.5 was chosen. Fluorescence changes related to spikes (red lines) were also subtracted. (**C**) Comparison of single-cell response curve without and with neuropil correction (r = 0.5). Error bars show sem. (**D**) Mean coefficient of variation of ΔF/F peak as a function of r, normalized to no neuropil subtraction (r = 0), for all mouse lines. Circles indicate individual cells.

**Figure 3-figure supplement 1.**
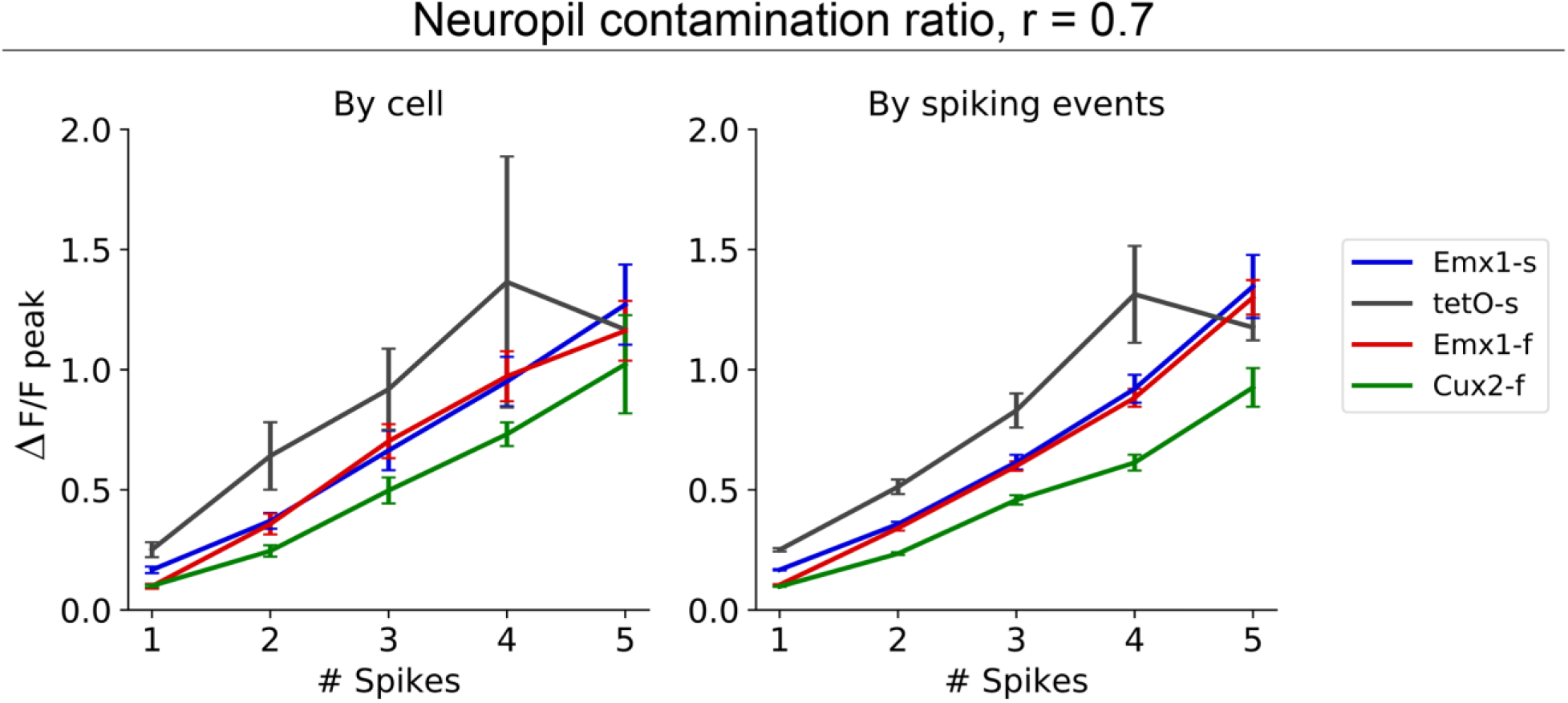
Population spike-to-calcium response curves with neuropil correction. Population spike-to-calcium fluorescence response curves computed from all spiking events (no exclusion of atypical spiking events) using a neuropil contamination ratio of 0.7. By cell: population response was computed from the mean response of individual cells (left). Emx1-s: n = 32 cells; tetO-s: n = 6; Emx1-f: n = 25; Cux2-f: n = 26. By spiking events: population response was computed from spiking events pooled from all cells as in Chen et al., 2013 (right). Emx1-s: n = 1264, 668, 298, 118, and 57 spiking events for 1-5 spikes; tetO-s: n = 254, 103, 48, 17, and 6; Emx1-f: n = 1300, 1010, 476, 183, and 70; Cux2-f: n = 2673, 788, 207, 118, and 46. Error bars show sem. Spike summation windows were 150 ms and 50 ms for GCaMP6s and GCaMP6f, respectively.

**Figure 4-figure supplement 1.**
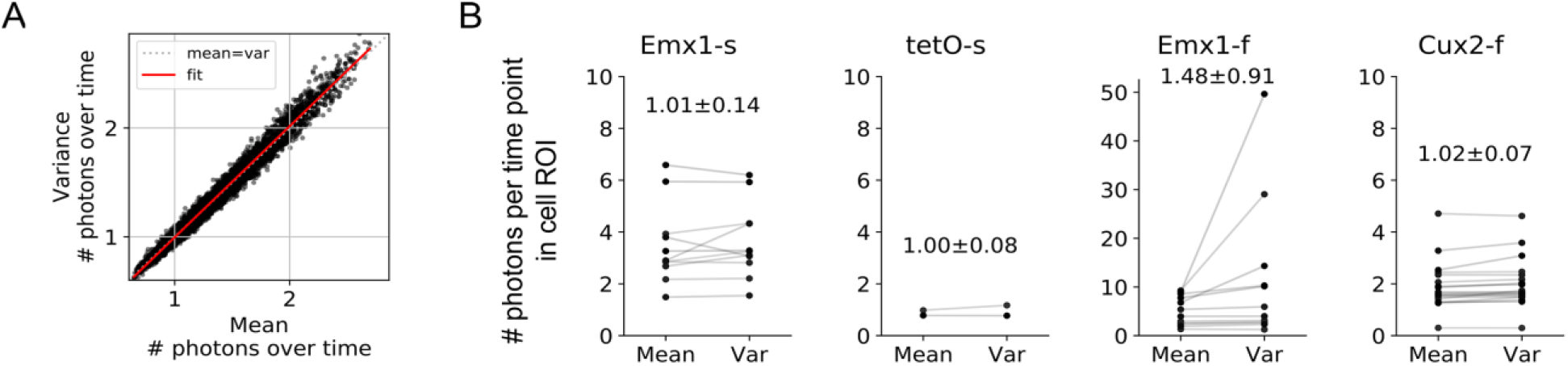
Relative contribution of shot noise to fluorescence response variability. (**A**) Relationship between variance and mean of the number of photons over time for all pixels in the FOV, for an example Cux2-f cell. (**B**) Mean and variance of the number of photons over time for pixels in the cell ROI in response to single-spike events. Fluorescence was converted to # photons at a time point corresponding to the maximum mean single-spike fluorescence response. Cells with ≥50 1-spike events were included in the analysis (Emx1-s, n = 11 cells; tetO-s, n = 2; Emx1-f, n = 12; Cux2-f, n = 19). Text labels show the variance-to-mean ratio (mean ± sd across cells). For each cell, variance-to-mean ratio was computed pixelwise for the cell ROI.

**Figure 4-figure supplement 2.**
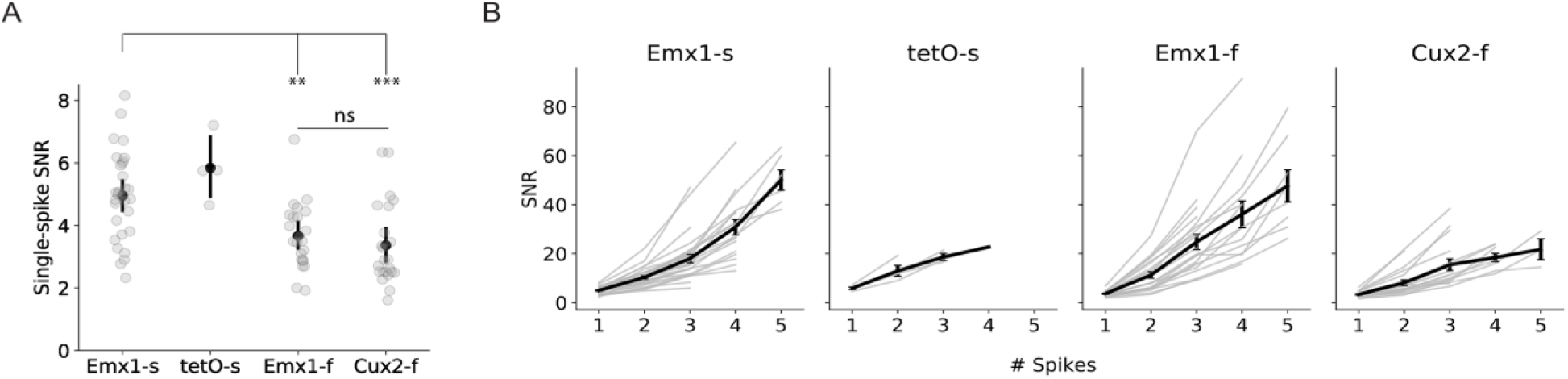
Signal-to-noise ratio. (**A**) Single-spike signal-to-noise ratio (SNR; single-spike ΔF/F peak normalized by standard deviation of ΔF/F during no-spike intervals; p = 4e-5, ANOVA; *, p < 0.05; **, p < 0.01, ***, p ≤ 0.001, multiple comparison test using Tukey’s honest significant difference criterion. Error bars show 95% confidence interval around the mean. (**B**) SNR as a function of the number of spikes. Error bars show sem. Spike summation windows were 150 ms and 50 ms for GCaMP6s and GCaMP6f, respectively.

**Figure 5-figure supplement 1.**
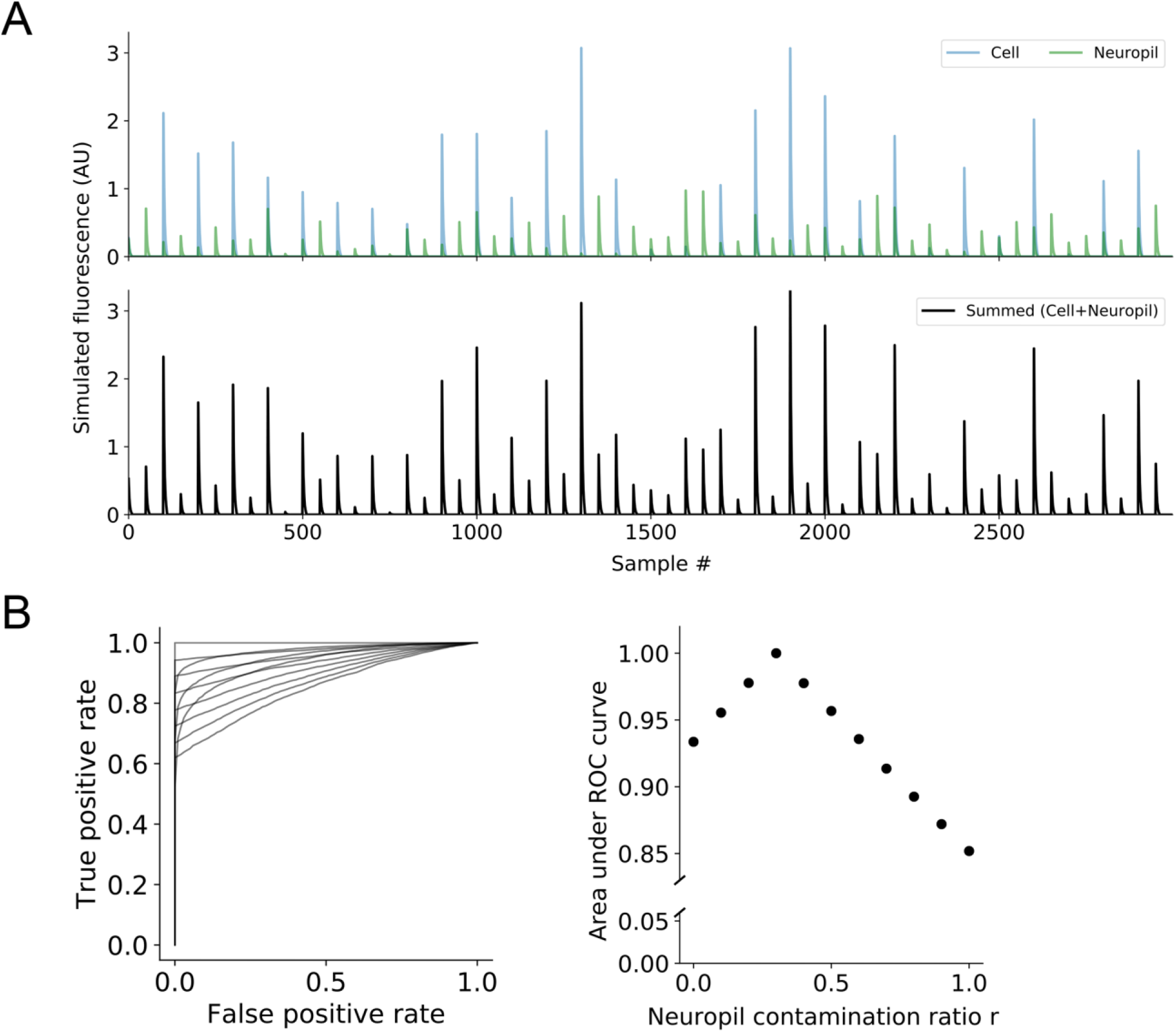
Simulated effect of neuropil subtraction on detection sensitivity. (**A**) Simulated cell and neuropil traces. The neuropil trace contained transients that were (1) associated with cell transients and (2) between cell transients, and amplitudes were scaled by the neuropil contamination ratio r (r = 0.3) relative to the cell amplitudes. (**B**) (Left) ROC curves for classifying cell amplitudes, where r was varied from 0 to 1 for neuropil correction. The detection threshold was defined as the x^th^ percentile of noise amplitudes (amplitudes of the neuropil trace that were between cell transients), where 1-x represented the false positive rate, and the detection rate (true positive rate) was the fraction of estimated cell amplitudes (amplitudes of the summed trace that were associated with cell transients) above the detection threshold. (Right) Area under ROC curves as a function of r.

**Figure 5-figure supplement 2.**
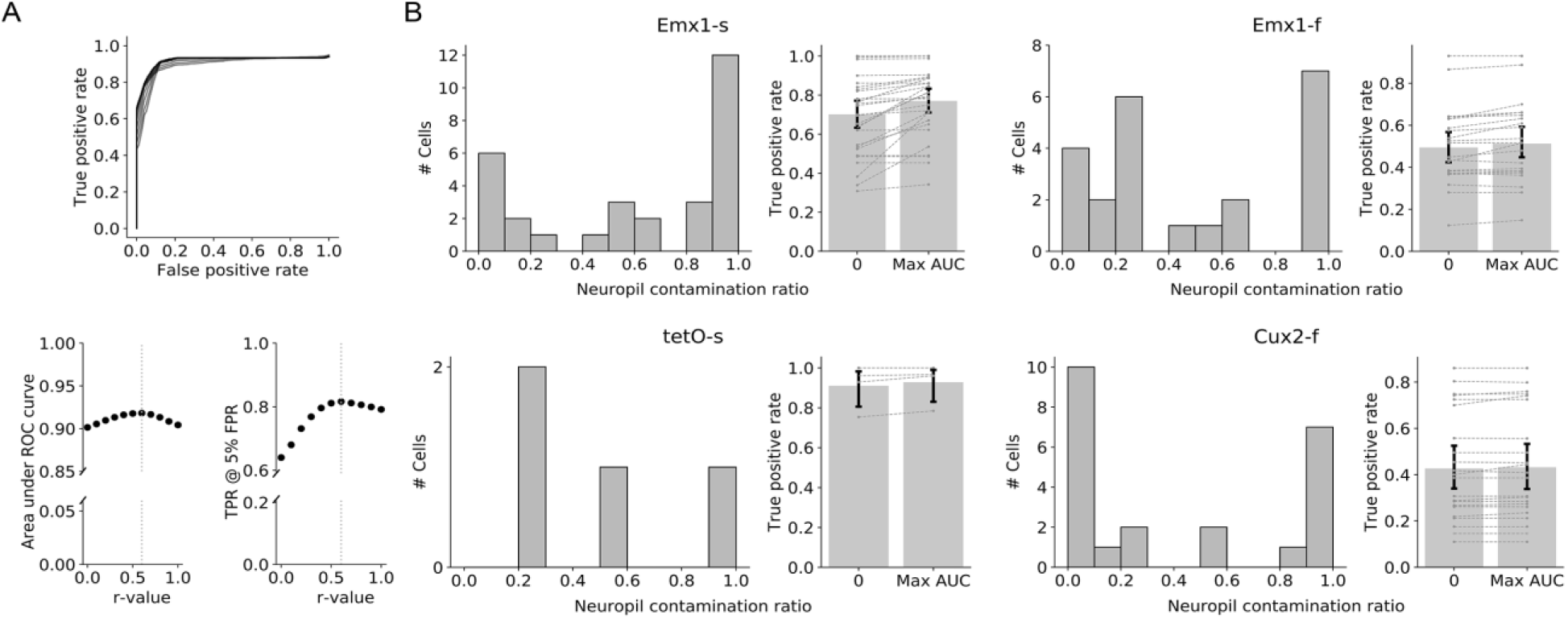
Effect of neuropil subtraction on single-spike detection rate. (**A**) Identifying the optimal neuropil contamination ratio r for maximizing spike detection efficiency. (Top) ROC curves for classifying single-spike events in an example Emx1-s cell, where the neuropil contamination ratio r was varied from 0 to 1. (Bottom) Area under ROC curve (left) and true positive rate at 5% false positive rate (right) as a function of r-value. (**B**) For each mouse line: (Left) Distribution of optimal r-values in individual cells for maximizing area under ROC curve. (Right) Single-spike detection at 5% false positive rate before (r = 0) and after optimization. Error bars show 95% confidence interval around the mean (Emx1-s: n = 30 cells with ≥1 no-spike interval; tetO-s: n = 4; Emx1-f: n = 23; Cux2-f: n = 23).

